# Cryo-EM structure and B-factor refinement with ensemble representation

**DOI:** 10.1101/2022.06.08.495259

**Authors:** Tristan Cragnolini, Joseph Beton, Maya Topf

## Abstract

Cryo-EM experiments produce images of macromolecular assemblies that are combined to produce three-dimensional density maps. It is common to fit atomic models of the contained molecules to interpret those maps, followed by a density-guided refinement. Here, we propose TEMPy-REFF, a novel method for atomic structure refinement in cryo-EM density maps. By representing the atomic positions as components of a mixture model, their variances as B-factors, and a model ensemble description, we significantly improve the fit to the map compared to what is currently achievable with state-of-the-art methods. We validate our method on a large benchmark of 366 cryo-EM maps from EMDB at 1.8-7.1Å resolution and their corresponding PDB assembly models. We also show that our approach can provide newly-modelled regions in EMDB deposited maps by combining it with AlphaFold-Multimer. Finally, our method provides a natural interpretation of maps into components, allowing us to accurately create composite maps.

## Introduction

Cryo-electron microscopy (cryo-EM) can resolve the structure of biomolecules at an ever-improving resolution. Larger complexes can now be visualised as 3-dimensional density maps at near-atomic resolutions, and in various conformations. The interpretation of those maps often hinges on fitting atomic models of the different macromolecules present in the complex^1–3^. This procedure is often difficult and requires the user to provide accurate models, and a well-estimated resolution (which can vary at different parts of the map). Pre-existing experimental or predicted atomic models may be in a different conformation and converging to a well-fitted one may require significant sampling.

Several methods are in common use for this procedure. To improve the map fit, the map can be treated as a scalar field, for which a gradient can be used as a force^4,5^. Optimisation of the position against the correlation coefficient (CCC) has also been proposed^6^, or by Bayesian expectation-maximisation against the density observed in the map^7,8^. The sampling itself is usually based on either molecular dynamics (MD)^4,9^, minimisation^10^, normal mode analysis and/or gradient following techniques^11,12^, or Fourier-space based methods^2^. Manual inspection and modification of the structure, or targeted sampling for specific parts of the structure, are also common, especially at high resolutions^13–15^.

Molecular dynamics-based refinement methods have the advantage of wider sampling but may result in locally distorted structures. This can usually be fixed by either clustering the resulting data^9^ or by minimising the structures at the end of the run^6^. The use of a forcefield (such as CHARMM^16^ or AMBER^17^) have the added benefit of ensuring that clashes are generally absent from the structure since they include parameterised van der Waals repulsion terms.

Virtually all methods rely on blurring the model (globally or locally)^18^ to compare against the experimental map, which poses an additional challenge for maps of flexible systems that will often exhibit significant resolution heterogeneity between flexible and rigid regions. This heterogeneity in the map can also result from adding up density maps from different reconstructions (e.g result of multibody or focused refinement) into a so-called “composite” map^19,20^. However, a systematic way to combine multiple maps into a composite map has not been proposed yet.

Flexibility is intrinsic to biomolecular systems, which presents a challenge for methods that tend to rely on a single structure representation. Methods using a population of models^21–25^ can provide an improved understanding of the fit between map and models^26^. Mixture modelling is a powerful framework to represent arbitrary density probabilities comprising several parts: by iteratively estimating the model parameters, and then re-computing the expected distribution, a (locally) optimal model can be generated^8^. We use this approach to estimate both the local spread of density around atomic positions and the background noise level.

Here, we propose TEMPy-REFF (REsponsibility-based Flexible-Fitting) -- an expectation-maximisation scheme using a mixture model that provides self-consistent estimates for the atomic positions and local B-factors (Fig. 1). The B-factor estimates converge over a wide range of starting values, with position refinement comparing favourably with state-of-the-art refinement methods, such as Phenix^27^. We show that the method can accurately treat maps with highly heterogeneous resolution. To assess the quality of the refined models, we have developed a new measure that estimates the resolvability of every residue and allows us to compare the fit of different parts of the model in regions of varying resolution. We demonstrate on a large dataset (from the CERES database http://cci.lbl.gov/ceres and additional cases), that our approach produces a significantly higher quality fit (higher CCC) to the density than what is currently achieved with some of the state-of-the-art methods, such as Phenix. Finally, we show how our method can be used for ensemble model generation, map segmentation and composition of multiple maps.

**Fig. 1.**
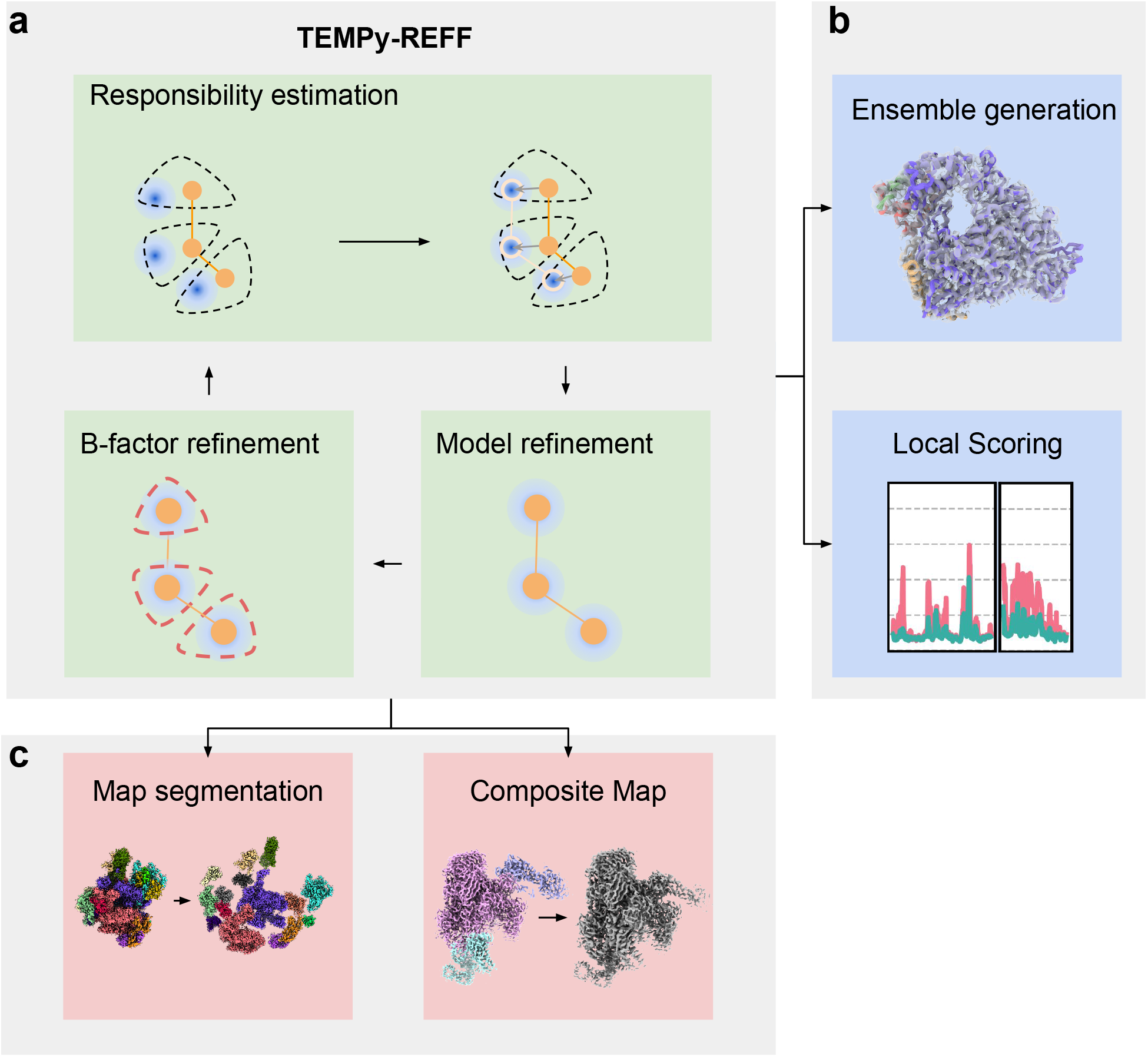
Flow chart summarising the steps in the TEMPy-REFF algorithm. **a**, The EM (Expectation-Maximisation) algorithm. Responsibility is an estimation of the part of the data that is represented by a given component in the mixture. New parameters (the mean and variance of each component corresponding to the position and B-factor value) for each component (e.g. for each atom) are then re-estimated using this responsibility and the experimental data. **b,** After refinement, an ensemble can be generated based on the local variance; local scoring provides a view of the quality of fit of all regions of the map, irrespective of the local resolution. **c,** By considering the sum responsibility of all the atoms in a chain, we obtain a natural expression of the part of a map represented by a given chain. This can be used for either segmentation or composition.

## Results

### Mixture modelling applied to refinement

We have developed a method based on Mixture Modelling (MM, using one Gaussian per atom and a uniform background term) to represent the estimated contribution of various parts of a model to the experimentally observed intensity. The Gaussians are fitted to the model in a self-consistent way, such that their summed contributions represent a (locally) optimal fit to the density. The intensity attributed to a Gaussian, or a sum of Gaussians, can be used to estimate their importance in representing a specific part of the map density. For example, by summing the Gaussians for atoms from a given protein chain, it is possible to determine which part of the map is best represented by this chain, or other chains, or are part of the general background noise in this map. Those weighted contributions (termed “responsibilities”) allow us to perform a variety of tasks that are commonly performed on cryo-EM maps (described in Fig. 1): fitting an atomic model to the map, segmenting the map into several parts, each representing a distinct entity (for example, a distinct subunit in a protein complex), or combining focused maps into a single overall composite map, with optimal weights of the focused maps.

Although Gaussian Mixture Model (GMM) approaches have been successfully employed before, this model was usually a simplified representation of the overall model and map^7,8,28^. By describing each atom as a Gaussian point spread function, a link between map and model is directly established: the intensity of each voxel is a direct sum of the contribution of each atom, as a function of its position, b-factor, and occupancy. It is important to note that the formalism used here does not require the use of Gaussian distributions, and alternative descriptions for the individual atomic contributions could be considered.

The responsibility calculation has several benefits: for regions of the map that are close to multiple parts of the structures, the mixture model allows for uncertainty in the assignment of the density. This “soft-mixing” improves the convergence of the refinement, by making it easier for structural elements to slide towards regions of density that are a better fit, even if they are currently fit to a high-density region of the map.

The calculation is also self-consistent, as is empirically demonstrated below: changes in the initial position and B-factor assignment for the structure result in identical or similar fit for a wide range of initial values.

### Benchmark B-factor refinement and assessment

The MM interpretation of the fitting problem allows us to obtain more accurate estimates of the B-factor that provide a locally optimal fit to the map (see Methods). While the B-factor optimisation is intended to be used together with position refinement, it is useful to test it independently, to assess the magnitude of the gain one can expect from it. This can be done by optimising the B-factors while keeping the atomic positions fixed. We performed such tests using our extended benchmark of 388 protein complexes (CERES+), which is based on the recently-released CERES^29^ database and additional cases (see Methods). The results are presented in Table S1.

The B-factor converges quite rapidly (Extended Data Fig. S1, S2), usually in 10-11 iterations or less, with minimal changes observed afterwards. This is due to the nature of the B-factor that describes the local spread of the data, which should remain similar for a given structure. The latter can be seen in the examples of Faba bean necrotic stunt virus (FBNSV) (EMD-10097) and the SARS-Cov-2 RNA-dependent RNA polymerase (EMD-30127) (Extended Data Fig. S1).

The B-factor assignment is highly robust, and the recovered B-factors are near-identical, after a few iterations (Extended Data Fig. S3, S4). Overall, they deviate by ±0.87 Å^2^ over 366 cases in our dataset. While additional changes in some B-factors can be observed as the structure itself is being updated, this is a feature of the change in coordinates (as the two are not independent) (Extended Data Fig. S5).

### Ensemble generation based on B-factors

To more accurately represent the variety of conformations that are compatible with the map, we generate a structural ensemble, based on the model we have obtained. We first generate models by randomly perturbing the positions of atoms, based on their B-factors, we then run a local l-BFGS^30^ minimisation (with openMM^31^), to locate close-by structures that are compatible with the data^32^. We then generate an ensemble map by averaging the blurred map obtained for all sampled structures in the ensemble.

The results can be seen in Fig. 2 for the refinement of rotavirus VP6 (EMD-6272) at 2.6 Å. The best-fitted model appears as only one solution among many. On the other hand, the ensemble average map exhibits a much higher quality-of-fit to the experimental map than any single model (Fig. 2a-b, Extended Data Fig. SS6). Interestingly, the local map-model agreement as measured by SMOCf correlates well with the difference in conformations of each residue between the different ensemble models (RMSFe, Fig. 2c).

**Fig. 2.**
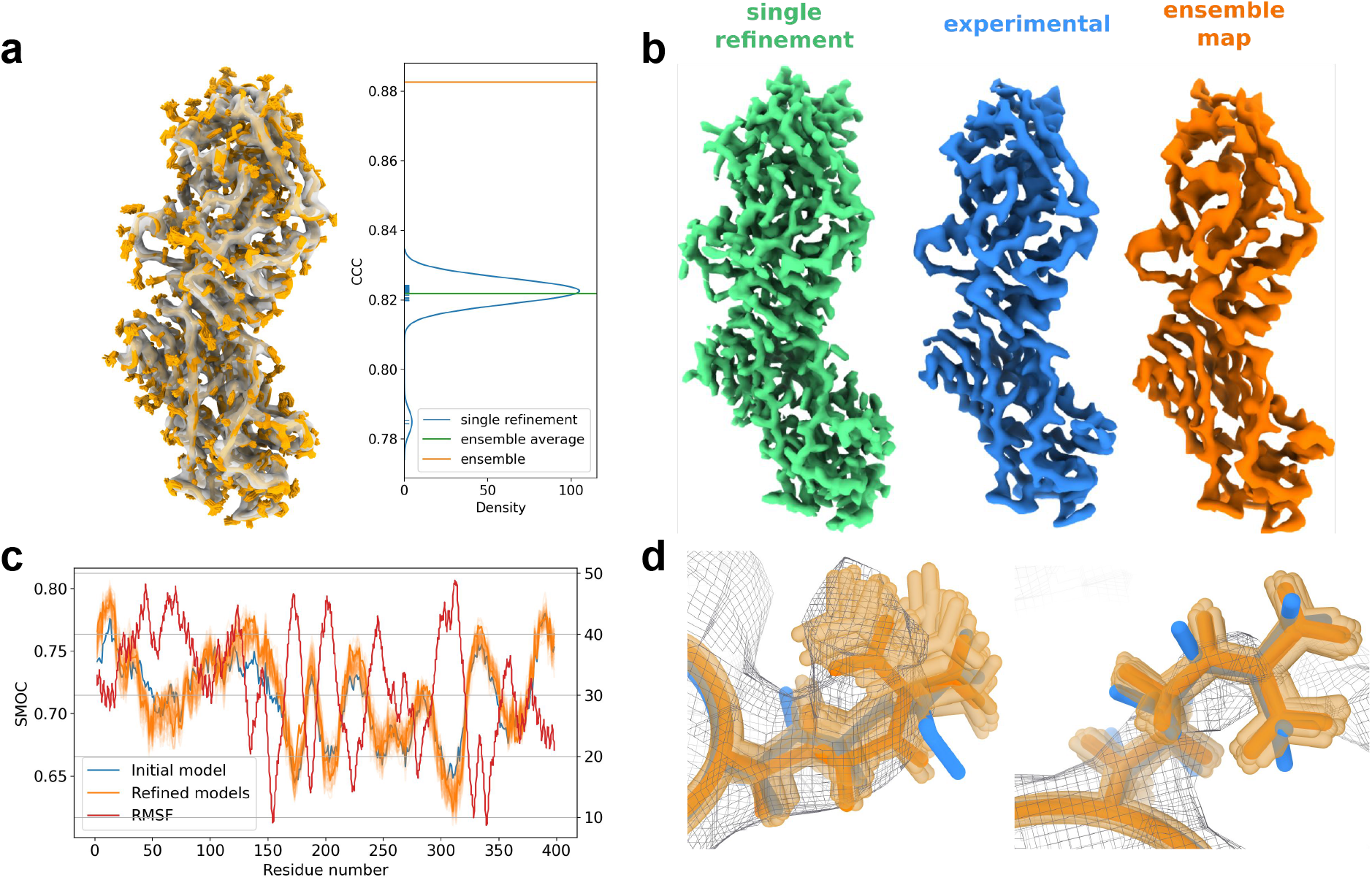
Ensemble representation illustrated on rotavirus VP6 (EMD-6272). **a,** Depiction of structure ensemble (orange), along with the map (transparent grey); a plot of the CCC of each individual model in the ensemble is shown (blue), as well as the ensemble map (orange); **b,** depiction of a single model map (green), experimental map, and our computed ensemble map at contour level 0.02; **c,** SMOCf plot and RMSFe for each structure in the ensemble; the RMSF and SMOCf score are clearly anticorrelated. **d,** Differences in the ensemble for different residues: for residue arg 107 (left) is more widespread, and the side-chain density is more spread out, only visible at a higher contour level, on the right side, for arg 217, the ensemble is highly constrained, and the side-chain density is well defined.

Intriguingly, the ensemble map resembles more closely the experimental map (Fig. 2a). The ensemble map density attributed to mobile side chains is spread out, while that attributed to more rigid ones is sharper (Fig. 2d). This would not be easily obtainable by simply simulating a map from a single given structure, even with the help of local B-factors. We determine the optimal number of models in an ensemble by calculating the CCC with the ensemble map generated from an increasing number of models (Extended Data Fig. S7).

### Benchmarking structure refinement

To assess the quality of improvement in the model using a large dataset, we look at the changes in average SMOCf^33^, CCC, MolProbity^34^, and CaBLAM^35^ scores between the initial conformation and after refinement. We benchmarked our method against the CERES dataset^29^ (see Methods), which is an automated Phenix^27^ re-refinement of 366 complexes in their cryo-EM maps at resolution <= 5 Å. The results are shown in Fig. 3 and Extended Data Table 2.

**Fig. 3.**
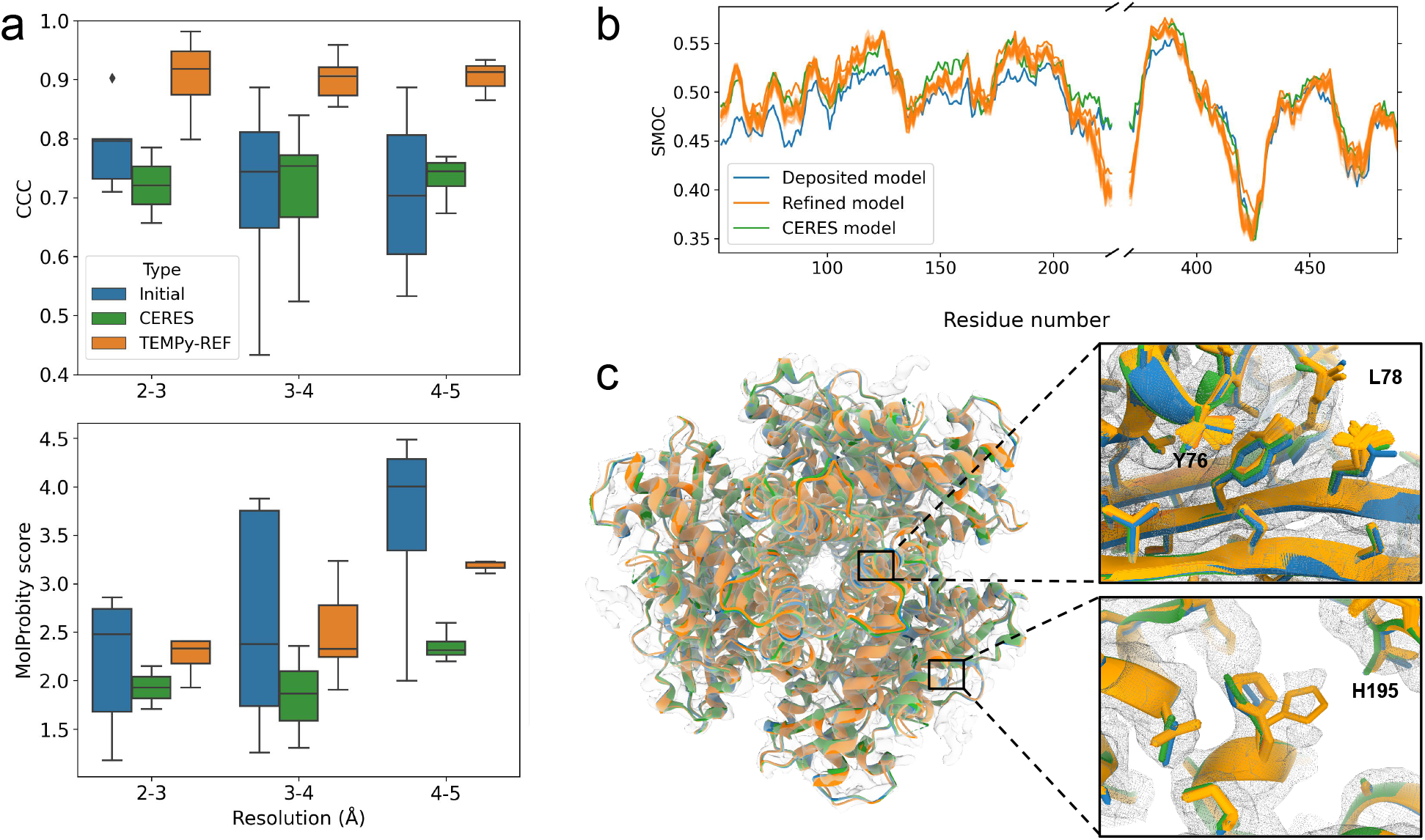
Refinement of the CERES benchmark and the example of the glutamate dehydrogenase refinement. **a,** Benchmark comparison using CCC, between the initial (PDB-deposited) models (blue), the CERES re-refined models (green), and TEMPy-REFF refinementbased model ensemble (orange), separated by resolution bands of 1Å. **b,** SMOCf plot calculated along the chain for the glutamate dehydrogenase deposited map and model (EMD-8194, 1.8Å, PDB ID: 5K12, blue), CERES-refined model (green) and the TEMPy-REFF model ensemble (orange): a higher score indicates a better local fit to the density for the corresponding residue. **c,** The deposited (blue), CERES-refined (green) and TEMPy-REFF ensemble (orange) models are shown in the map (grey mesh); insets of specific regions with improved quality-of-fit (around residues H195, Y76, L78) are shown.

Generally, with TEMPy-REFF we obtain very large improvements in CCC scores even for models that already have a good initial fit to the density, up to around 0.9 (close to the theoretical maximum of 1, which would indicate identical simulated and experimental maps). The CCC scores stay constantly high across resolution bands (Fig. 3a), while they become lower for the deposited and the CERES models. These improvements are statistically significant compared with results obtained for the deposited and the CERES models using paired-sample T-tests, the average improvements are 0.197 with a p-value of 0.0006 compared to EMDB models and 0.150 with a p-value of 0.003 compared to CERES models. Our MolProbity and CaBLAM scores are in line with deposited models. Interestingly, the MolProbity scores obtained with CERES stay in a very narrow range, while we see higher variability with TEMPy-REFF (Fig. 3a). This trend also holds true for CaBLAM scores (Extended Data Table 2).

Overall local measures of fit quality (SMOC) show improvements with TEMPy-REFF models. In the example of glutamate dehydrogenase (PDB ID 5K12, EMD-8194), this is apparent in most regions of the modelled structure (Fig 3b). A visual comparison between the initial (PDB-deposited) model, CERES-refined model, and TEMPy-REFF refinement-based model ensemble is shown in Fig 3c: while all structures are well fit to the map overall, insets of residue fit show the source of improvements: a histidine (H195) that is present in the map in two alternative conformations, with structures in the ensemble adopting either (bottom inset), or uncertainty in the exact side chain conformation (bottom inset) of two residues (Y76 and L78).

### Segmentation and composition

To tackle the problem of segmenting a map of a system comprised of multiple subunits, we use a well-fitted model, combined with our MM representation, to assign the intensity of each voxel in the “parent” map to different “child” maps: for each voxel, nearby models are assigned a portion of the voxel’s intensity proportional to their relative assigned responsibility for that voxel (see Methods). We demonstrate this on the 2.8 Å resolution map of glycoprotein B-neutralizing antibody Complex from Herpes Simplex Virus 1 (EMD-21247^36^) (Fig. 4a). To compose maps together, we follow the opposite approach: we assume that the maps are components of a parent map. We reconstruct this parent map, using the normalised intensities of each “child” map, and combine them to form the parent map, as demonstrated in Fig. 4b on RNA polymerase II super elongation complex (EMD-12966 to EMD-12969^37^).

**Fig. 4.**
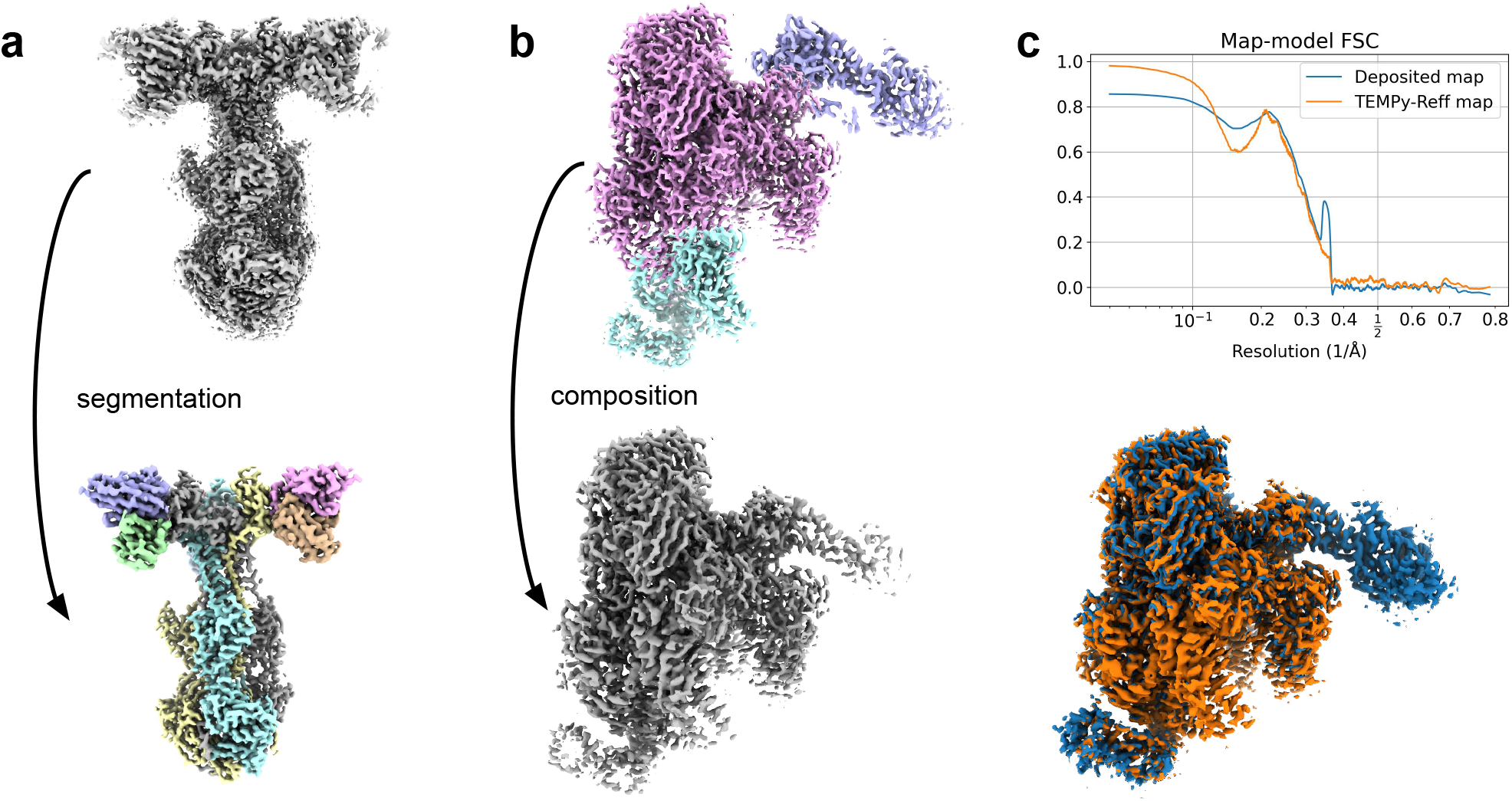
Using TEMPy-REFF for map segmentation and composition. **a,** Segmentation of maps (glycoprotein B-Neutralizing Antibody Complex from Herpes Simplex Virus 1, EMD-21247^36^) into 9 submaps corresponding to single chains. **b,** Composition of 3 focused maps into a composite map, using EMD-12966 (cyan), EMD-12967 (purple), and EMD-12968 (pink). **c.** Top: Map-model FSC between the deposited model and the deposited composite map, EMD-12969^37^ (blue) and between the deposited model and TEMPy-REFF composite map (orange). Bottom: TEMPy-REFF composite map (orange) superposed on the deposited composite map (EMD-12969, blue).

In both examples, this approach has several advantages: because the responsibility decays smoothly, there are no “seams” between segmented maps, or within composite maps, as evidenced in the FSC curve (Fig. 4c). Additionally, areas where the assignment would be uncertain are treated as such, and the density will not be arbitrarily assigned to a specific model or submap.

### **Case study 1:** yeast RNA polymerase III elongation complex

We explore the effectiveness of the TEMPy-REFF approach in more detail by refining the model of yeast RNA polymerase III elongation complex (PDB ID 5FJ8). The corresponding cryo-EM map (EMD-3178) was resolved at a global resolution of 3.9 Å^38^. A brief observation of the deposited model (which is hitherto referred to as the “deposited model”) suggests that it is well fitted to the cryo-EM map: we computed a position-optimised correlation using the ChimeraX^39^ *fitmap* utility, which gave a model-map correlation score of 0.82. The validation statistics presented in the PDB are reasonable; clash score of 14, Ramachandran outliers 1.1% and sidechain outliers 2.1%, with an overall MolProbity score of 2.8. Altogether, these observations make PDB ID 5FJ8 an interesting test case for the automated refinement using the TEMPy-REFF procedure as it allows us to measure our automated algorithm against a manually optimised model-building pipeline that produced a good quality model for a map with heterogeneous resolution.

As we require slight changes to the model (namely the addition of hydrogens), we also compute the correlation of this prepared model, which is also 0.82, showing no significant change compared to the initial model. The ensemble models resulting from TEMPy-REFF refinement show a significant improvement in the correlation to the map, with a final CCC of 0.94. This is in large part due to improvements afforded by the ensemble representation, with some improvements at the single-model stage due to better positioning of atoms. (the final model CCC, without the ensemble, is 0.83). A representation of the model, as well as the quality-of-fit for each chain, is shown in Fig. 5.

**Fig. 5.**
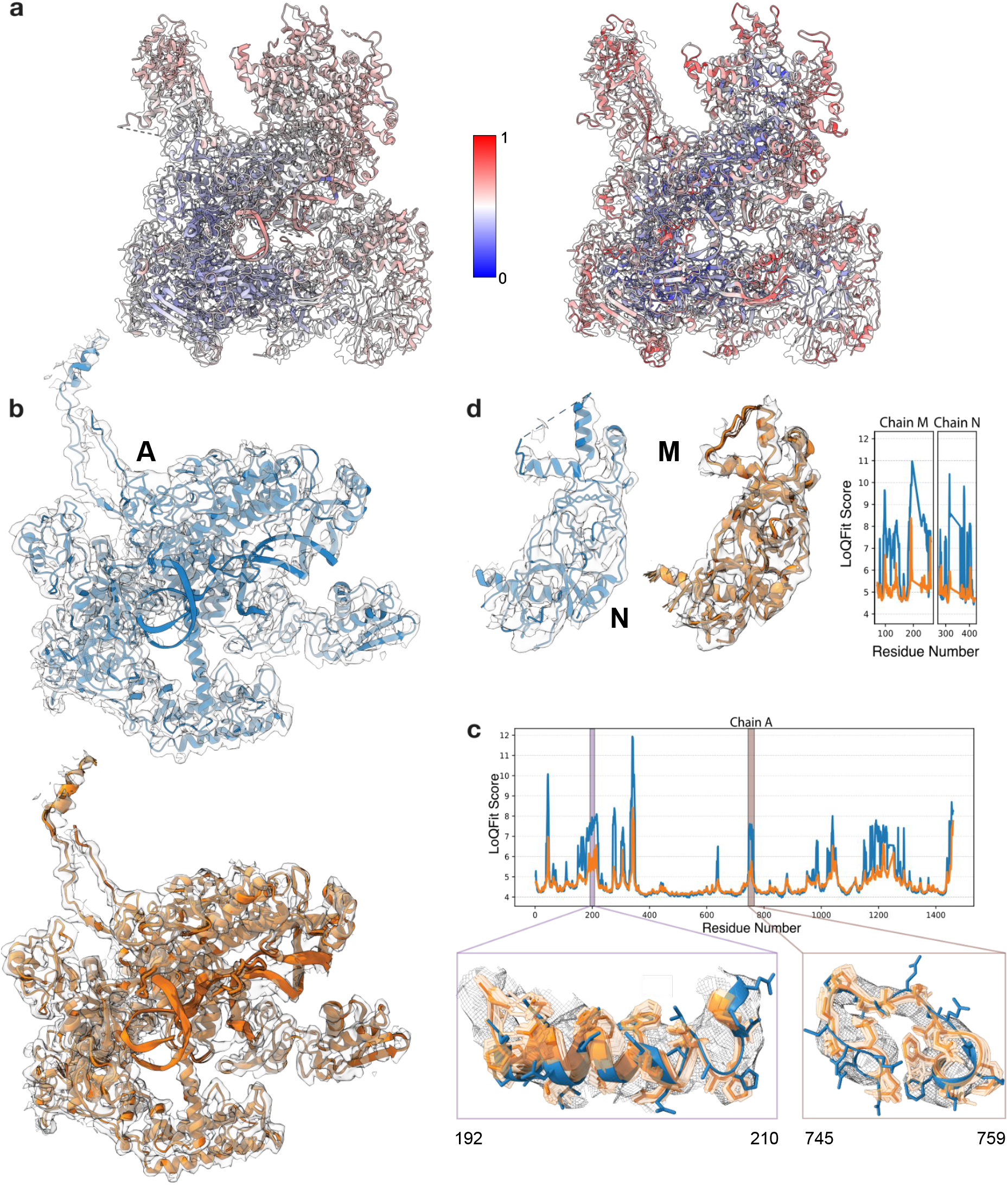
Case study of RNA polymerase III elongation complex. **a,** The deposited model of the RNA polymerase III elongation complex (PDB ID 5FJ8, left) and the TEMPY-REFF refined model (right) coloured according to the B-factors (calculated with RELION (left) and TEMPy-REFF (right), and the 3.9 Å resolution cryo-EM map (EMD-3178) (transparent grey). For comparison purposes, the B-factors are normalised between 0-1. **b,** Chain A in complex with RNA (chains R, S and T), from the deposited structure (blue, top), and the refined model (orange, bottom), both in the map (grey mesh). **c,** LoQFit scoring of chain A before (blue) and after (orange) refinement with TEMPy-REFF. Insets show several regions before and after refinement (residues 192-210, 745-759) colored as in **b,**) and the ensemble of models (lighter orange). **d,** Chains M and N, before and after refinement (left), with the change in LoQFit scoring (right) (colored as in **b,**).

We next apply the TEMPy LoQFit score (see Methods) to locally assess the improvement of our TEMPy-REFF refined model, versus the deposited model. We visualise the LoQFit score at each residue in both models using 2D plots (Fig. 5). The average LoQFit score for the deposited model was 10.1 Å, and model agreement was particularly high in chains A and B at the central regions of the model and map, where the average LoQFit score was 5.4 and 5.6 Å, respectively. However, even in these regions we observe peaks in the LoQFit score, consistent with poorer model fit, such as those seen around residues 192-210 and 745-759 in chain A (Fig. 5c), as reflected in the higher B-factors in this region (Fig. 5a). In addition to this, we identify extended regions of poorer model fit, generally occurring within chains that lay at the edge of the complex in solvent-exposed regions with poorer resolution, including chains M and N (Fig. 5d). In these chains the average LoQFit score was 7.4 and 7.0Å, respectively, reflecting the lower map resolution (and correlating with high B-factors), as well as poorer model fit in the deposited model. Refinement with TEMPy-REFF resolved many of these poorer fitting regions: the average LoQFit score for the refined model improved to 9.0 Å, and we observed significantly better model fit at lower resolution regions of the map. The average LoQFit score for chain O improved to 5.9 Å in the refined model (from 7.3 Å, data not shown), and in chains M and N the average LoQFit score improved to 6.0 and 5.7 Å after refinement.

### Case study II: Nucleosome-CHD4 complex structure

The nucleosome is a large nucleoprotein present in the nucleus, which is the primary effector in the compaction of DNA. High-quality reconstructions have been obtained, but its dynamic nature and strained DNA strands wound around the histone proteins make it a challenging system to obtain a good structural model. We apply TEMPy-REFF to refine the model associated with map EMD-10058^40^ (PDB ID 6RYR) (Fig. 6a-d). The deposited cryo-EM map clearly suffers from very variable resolution (range: 3-10 Å, see Extended Data Fig. S8) which affected the quality-of-fit of the deposited model (Fig 6a). Following refinement, the local details of the map are well respected, especially showing improvement in the DNA structure, as reflected by the SMOCf score (chain I and J, Fig. 6c). Nucleic acids are often present in biomolecular complexes resolved by Cryo-EM, and refining their geometries with respect to the map is an important part of model refinement. In the deposited model, local deformations pull the bases slightly away from the density, and from the expected geometries to allow hydrogen bond formation. Our automated refinement pulls them back, forming hydrogen bonds in the process (Fig. 6d). The LoQFit and the local resolution follow similar trends (Extended Data Fig. S8).

**Fig. 6.**
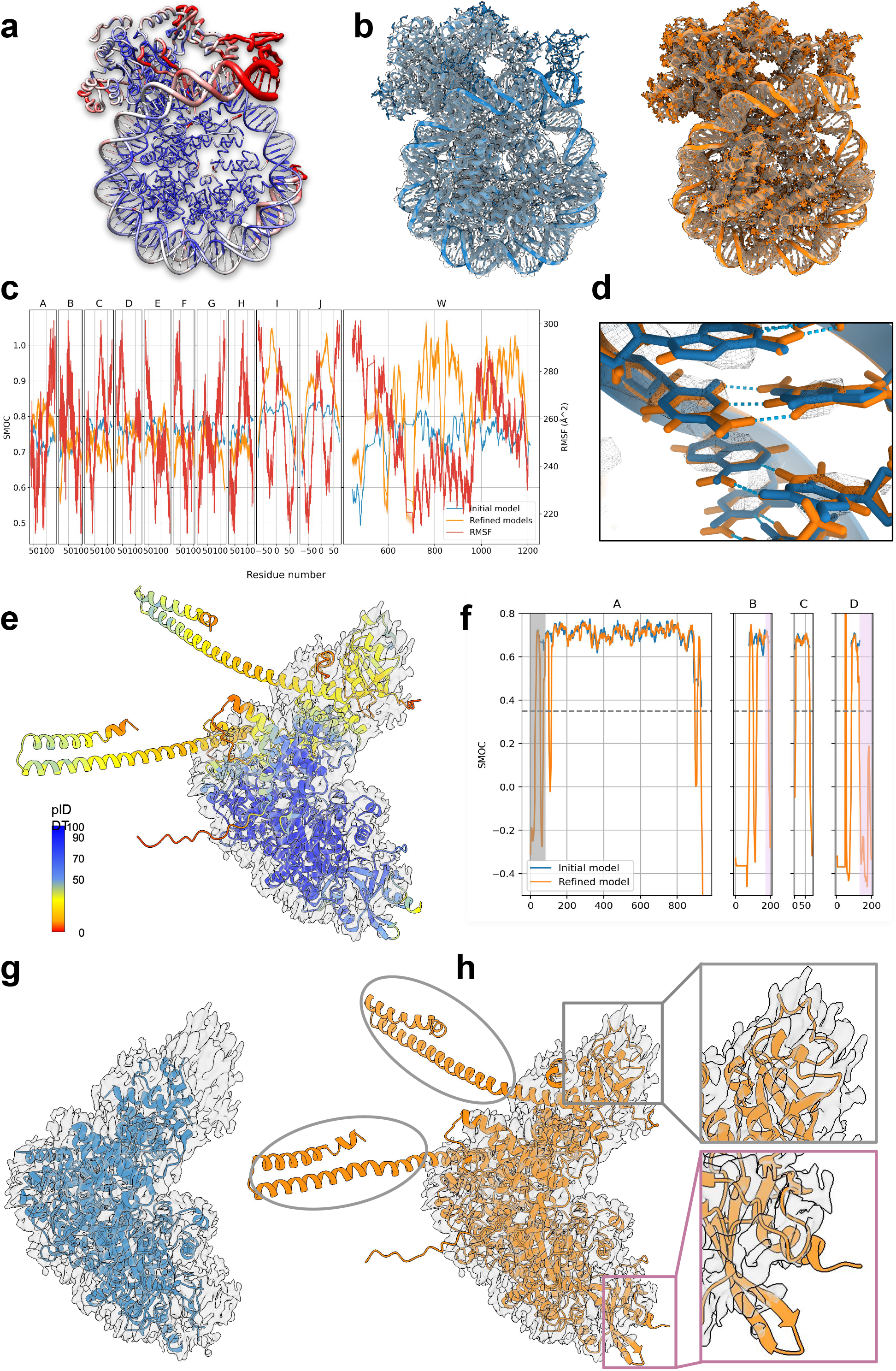
Case studies of Nucleosome-CHD4 complex and SARS-CoV-2 RNA polymerase (AlphaFold model refinement). **a,** A nucleosome structure in complex with chromatin remodelling enzyme CHD4 (EMD-10058, PDB ID: 6RYR) is shown (worm representation), with the width proportional to the B-factor, and color based on local resolution (computed with ResMap). **b,** Deposited model (left, blue) and the ensemble of models and ensemble map calculated with TEMPy-REFF (right, orange), shown inside the cryo-EM map (transparent grey). **c,** SMOCf and RMSFe plot for each chain. **d,** Zoom-in on some of the DNA basepairs (chain I/J, basepair 54) fitted in the map (mesh representation). The deposited model is shown in blue, TEMPy-REFF model in orange and hydrogen bonds are indicated in cyan. **e,** AlphaFold2 predicted structure, with the colouring indicating the plDDT confidence measure (blue means higher confidence, red means lower confidence), fitted in the deposited map (EMD-30127, grey) **f,** SMOCf plot of the deposited and refined model (blue before refinement and orange after). The regions which are highlighted in grey and pink (correspond to inset regions in Fig. 6h) contain residues which are not present in the deposited model but are present in the AlphaFold2 model and are well fitted to the map. **g,** Deposited model for the SARS-CoV-2 RNA polymerase (PDB ID 6M71, blue) fitted in the deposited map (transparent grey). Unassigned regions are visible, top and bottom right of the map. **h,** TEMPy-REFF model (orange) obtained by refining the AlphaFold2 prediction in the deposited map (transparent grey). Newly modelled regions that fit in the density (as in Fig. 6h) are shown with coloured squares. The regions that are circled do not correspond to density and have low plDDT and SMOCf scores.

This case study also further demonstrates how the ensemble map calculated with TEMPy-REFF has greater similarity with the experimental map than a single model (either the deposited model or a single-refined model). Specifically, the top domain, which shows greater variability (also reflected in the worse local resolution in this region, Fig. 6a), is also more variable (higher RMSFe) in our ensemble map (Fig. 6b-c).

### Case study III: SARS-CoV2 RNA polymerase and AlphaFold2

To refine a model into an experimental cryo-EM map, an initial model is needed. Although building a reliable model directly from the map is sometimes possible, in most cases this cannot be done reliably as the resolution is not sufficient to allow a reliable assignment of every atomic position. It is more common to start by creating a model based on structural homologs of the system of study. For systems with limited structural homologs or none, an initial model may be obtained by deep-learning based *ab initio* tools, such as AlphaFold2^41^ or RosettaFold^42^. Nowadays, such programs are able to create very high-quality protein models^43^. The predicted lDDT score^41,44^ (plDDT) is also an excellent tool to decide which part of the model can be reliably kept, and which may not be correctly predicted, due to flexibility or lack of known homologous sequences and structures.

To assess the capability of our method to refine such a model, we used AlphaFold2 to create a model of the SARS-Cov-2 polymerase. We used the polymerase sequence (UNIPROT ID: P0DTD1, residues 4393-5324) and only considered templates present in the PDB at least a year earlier than the deposition date of the deposited model (PDB ID 6M71)^45^. The predicted model was refined into the SARS-Cov-2 polymerase cryo-EM map at 2.9 Å resolution (EMD-30127) (Fig 6e-f). The resulting model (Fig. 6h) is highly similar to the deposited model (Fig. 6g) at most residue positions, which was calculated using Chimera^46^, Coot^14^, and Phenix^27^. However, more intriguingly, using a SMOCf plot, we show that some residues that were not present in the deposited structure^45^ can actually be placed into the map, with fitting scores much greater than chance (Fig. 6f and h).

## Discussion

We have presented TEMPy-REFF, a novel structure refinement protocol that takes into account the local features of a cryo-EM map using a mixture model with an error term. Our approach naturally incorporates both position and B-factor estimations in the same framework. This information is essential to represent the local variability around atomic positions.

Currently one of the greatest challenges in model building into cryo-EM maps is, how we can evaluate the quality-of-fit in a system not described by a single resolution value, but rather varying “local” resolution. We address this challenge using B-factor estimation. We find, as previously shown by us and others^21–23,25^, that an ensemble of equally-well fitted models represents this local variability better than a single model. However, we go one step further, by showing that an “ensemble map” calculated from these models, provides a better representation of the experimental map, in comparison to a “traditional” simulated map (which is typically generated from a single Gaussian function per atom) (Fig. 2a). This is best showcased in Fig 3c, where a double occupancy site for a histidine necessarily requires more than one model to be correctly represented. The improvement is also evident in regions of lower local resolution (Fig. 6a,c), which may indicate an inherent local flexibility of the structure, although this cannot be easily deconvolved from the blurring due to optical factors^47^, or the 3D reconstruction procedures.

Ensemble methods have been common practice in the NMR community and have been suggested as a way of dealing with the uncertainty in the data^22,23,32^. This has also been demonstrated previously for X-ray crystallographic data^48^, and we similarly observe a plateau as more models are added to the ensemble (Extended Data Fig. S7). Furthermore, when analysing the differences on a local level (for example at the residue level) using a distance measure (such as the RMSFe), we observe that the local-fit-quality (using SMOCf) correlates well with those differences (Fig. 2c). As demonstrated on our new local scoring method and unsurprisingly, the local-fit-quality also correlates with the local resolution estimation (such as ResMap^49^).

Overall, our automated refinement procedure is computationally quick (Extended Data Table 3), resulting in high-quality models that are well-fitted to the cryo-EM map and with good model quality as assessed by MolProbity. Without the ensemble representation of the fitted models, the local and global model-map fit score is comparable or higher than Phenix on average (as represented by the 366 CERES results). It is worth noting that the quality-of-fit of our ensemble procedure remains near-constant at lower resolutions, while the quality-of-fit decreases for CERES, as well as for our single refinements. This shows again that the ensemble approach captures an important aspect of the model-map fit, which is missing from single model refinement approaches.

The change in MolProbity and CaBLAM scores in the refined models exhibit less spread after refinement: models with a low number of structural outliers (as defined in CaBLAM) tend to increase slightly, while models with high numbers of outliers tend to decrease significantly. However, this effect is much more pronounced in Phenix-refined models, possibly due application of stronger “structural” restraints. On one hand, this may be a desirable behaviour, as outliers should be uncommon, but on the other hand, they should be nonetheless present, if supported by the data. Furthermore, a lower number of outliers does not guarantee a better model.^50^

Since 2018, deposition of composite maps has been increasing significantly due to a growing number of macromolecular assemblies for which focused maps for different assembly subunits are obtained (often due to conformational flexibility). Some methods have been proposed to compose such maps^20^, however, there is currently no systematic way to evaluate this. Here, we provide a self-consistent way to perform this procedure. Our approach has the advantage that the responsibility decays smoothly, i.e., there are no “seams” between segmented maps, or within composite maps: areas where the assignment would be uncertain are treated as such. The same holds for segmentation, since the density will not be arbitrarily assigned to a specific model. However, the method also has some drawbacks, the clearest of which is that errors in modelling will result in errors in segmentation or composition. This problem can be remediated by refining the model in the map (or at least assessing the quality-of-fit).

Finally, we show that our refinement protocol can take advantage of recent developments in the field of protein structure prediction^41,42^. Starting refinements from AlphaFold2^41,42^ models is not only possible, it gives results on par with manual refinement (despite using an automated procedure) and highlights that better and more complete models can be obtained by using our automated refinement approach, including more residues that are sustained by the map information (Fig. 6 eh). This paves the way for more reliable, and reproducible model building and assessment, with an automated procedure from start to finish, where alterations in refinement protocols can be objectively and continuously assessed ^43,51^.

Further work will be needed to understand the impact of ensemble model representation, and how to use such an approach in assessing model-map fit quality, especially for inherently flexible protein assemblies observed by cryo-EM. The ensemble can be thought of as sampled from an underlying, continuous distribution, and more work may be needed to accurately represent this distribution. Large scale exploration with molecular dynamics could be a suitable tool to provide better resolved conformational ensembles.

## Methods

### Definitions

Given a reference experimental 3D map *M_E_*, with an estimated resolution R, we will attempt to improve the fit of a model comprised of N atoms, that has estimated coordinates 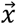, B-factors 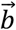, and occupancies 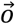 (which we will set to 1 and ignore thereafter).

### Algorithm

The full algorithm can be summarised below with detailed explanation following:

- Initialise coordinates 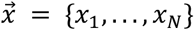, B-factors 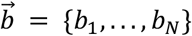, and resolution *r*.
- Loop over:

- Compute conformational energy and forces 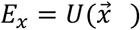 and 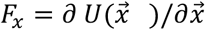, for a given forcefield.
- Compute a simulated density map 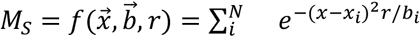 Compute map-derived forces 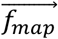.
- Change coordinates *X* using 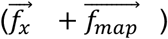.
- Change B-factors and occupancies based on the experimental cryo-EM map, *M_E_*.

### Simulated cryo-EM map

From the position X and the B-factors B, we can compute a simulated map, *M_S_*, by adding up the expected intensity due to each atom in a given group of atoms, modelled with a Gaussian distribution:

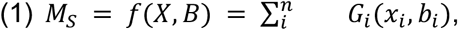

with each Gaussian taking the form:

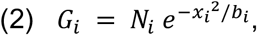

with *N_i_*the atomic number of the corresponding atom with index *i*, *x_i_*the cartesian coordinates, and *B_i_*the B-factor. Although it is possible to define a 3-dimensional B-factor, we restrict ourselves to a scalar for our refinement.

Finally, we consider the error (which comprises background noise, reconstruction errors, and additional components) as a uniform distribution across the whole map, determined by a single scalar parameter:

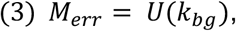

with *k_bg_* the uniform distribution parameter, corresponding to the average background noise level. We thus obtain the final expression for the simulated map as follows:

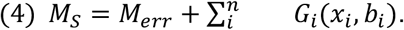

And re-estimation of k_bg_ is done using

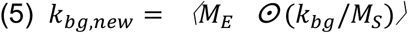

where *M_E_* and *M_S_* are the reference map and experimental map, respectively.

### Conformation-based force calculation and molecular dynamics (MD)

OpenMM is used for the conformation-based force calculation and molecular dynamics^31^. We tested CHARMM36 and AMBER in OpenMM (Extended Data Table S4), and they show slight differences in the preferred backbone dihedrals (Extended Data Fig. S9). Although other forcefields are available, we use AMBER for our runs. We use the GB-Neck2 implicit solvent model^52^, with κ=3.

### Map derived force calculation and B-factor calculation

We consider two methods to improve the fit quality: MD, where the system’s coordinates are integrated over time, taking into account the forces atoms exert on each other; and energy minimisation, where the coordinates of the system are changed to minimise a function.

To combine our description of the map with the energy terms that are usually present in forcefields, we compute a fictitious force representing the direction of the change in position induced by the Gaussian fitting (for MD), that can be expressed as *F_map_* = *K δ*(*x_i_*), with K = 400 a constant, and *δ*(*x_i_*) the estimated change in position shown below.

The quantities associated with each atom *i* (x_*i*_, b_*i*_, o_*i*_) are dependent on an estimated experimental density attached to a given atom, which can be expressed as follow:

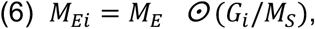

with the reference map *M_E_* and simulated map *M_S_* (as described above), and *G_i_*, the Gaussian generated by atom *i*. This can be thought of as a weighted average based on the reference map, and the weighing factor *G_i_* / *M_S_* is known as the responsibility in mixture modelling.

The new position of atom *i* is estimated as the real-space coordinate average (centre of mass) in the intensity-weighted map for atom *i, M_Ei_*:

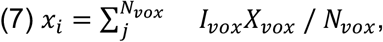

with a sum over the intensity *I_vox_* in position *X_vox_* of each voxel, and *N_vox_* the total number of voxels.

The B-factor is interpreted as the variance of the estimated distribution *M_E_i__*:

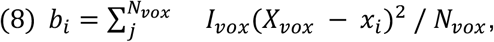

with *x_i_* the re-estimated position above.

Although occupancies could be computed independently for each atom, in a similar manner to the position and B-factor, it is expected that nearly all atoms in a chain should be present at a similar frequency. Therefore, we set the occupancies to 1, which simplifies the optimisation procedure, by reducing the number of free parameters.

### Generation of segmented and composite maps

Segmenting a map into several sub-parts is a common post-processing task, useful to both visualise and further analyse the obtained 3D map. A well-defined procedure for this task can be obtained from the mixture modelling formulation we provide, by noting that the responsibilities attributed to each chain of a model can be used to weight the intensities of the map. In this way, we can obtain a map corresponding to a specific part of the model, for example corresponding to a specific chain *j*:

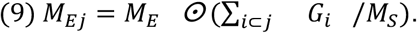

This formulation also allows us to combine multiple maps reconstructed from the same dataset, obtained for example with focused or multibody refinement protocols^19^. Normal addition or averaging will incorrectly represent the transition regions on those maps (see Extended Data Fig. S10), since the boundaries between the reconstructed subvolumes, present in each map, are usually not sharp. By weighting the intensity in each map, we smoothly transition between their estimated intensities, without over or undercounting the intensity in boundary regions (Extended Data Fig. S10):

For multiple experimental maps M_i_, we obtain a composite map:

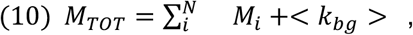

Where *N* is the number of maps and < *k_bg_* > is the average uniform distribution from all maps:

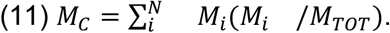

### Local quality of fit (LoQFit)

We implemented a new local fit quality score as part of the TEMPy2 python package. The new score – LoQFit – uses an approach similar to a local FSC score for cryo-EM maps ^53^ in order to assess the fit quality of a protein model. This local FSC score is calculated for regions defined by a soft-edged spherical mask, centred at the C_α_ atom for each residue in the fitted model and applied to both *M_S_* and *M_E_*. The diameter of this mask is 5 x the global resolution of the experimental map. We use an FSC threshold of 0.5 to determine the LoQFit score for each residue. To improve the smoothness of the final LoQFit plot, we include an option to estimate the exact frequency at 0.5 correlation between the two maps, using linear interpolation.

We also use SMOCf to estimate the local quality of fit^33^. Briefly, SMOCf uses a local window around each residue, and then computes the Manders overlap coefficient between the simulated observed maps in this region.

### Ensemble algorithm

To compute an ensemble representation, we create an ensemble of locally perturbed conformations, with displacement sampled from a Gaussian with a variance equal to the B-factor. We then locally minimise each model in the ensemble, to keep acceptable stereochemistry. We then generate blurred maps for each conformation in the ensemble, and compute a voxel-based average. To determine the number of models in an ensemble we increase the number of models until there is no increase in CCC. This average blurred map represents the final ensemble average map we use throughout the text.

### RMSFe

To compute the RMSF value for our generated ensemble, we first compute the mean structure, and then compute the RMSF using the normal formula. For an ensemble of structures with position matrices {r_1_, r_2_ r_N_}:

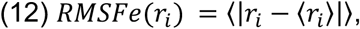

with the brackets indicating an average, and the bars a euclidean distance. To have one RMSFe value per residue, our position matrix *r* contains only C_α_ positions

### Local resolution calculations

We used the ResMap method to compute local resolution estimates^49^. ResMap uses local windows of varying size, and statistical tests to determine the most likely resolution for each voxel in the map.

### Generation of benchmark and assessment

Our benchmark is based on the CERES database^29^ (Extended Data Table 2). We took the corresponding deposited maps and structures from EMDB^54^ and PDB^55^, and the re-refined structures from CERES. Because of the CERES database setup, our benchmark contains maps resolved from 2 to 5 Å resolution. After our own refinement, we compare the CCC using the command line Chimera interface for *fitmap^39^*, molprobity and clash score using *phenix.molprobity*, and CaBLAM using *phenix.cablam^56^*.

## Acknowledgements

We thank Tom Mulvaney, Dr. Aaron Sweeney and Dr. Sony Malhotra for helpful discussions and Dave Houldershaw for computer support. Tristan Cragnolini and Maya Topf were supported by the Wellcome Trust grants 209250/Z/17/Z.

## Extended Data

**Extended Data Table S1.** Change in CCC upon B-factor refinement in TEMPy-REFF. Initial values are listed for a uniform B-factor of 10, final are listed after 5 B-factor refinement steps. Atomic coordinates are *not changed* during this refinement, as this is to test the effect of B-factor refinement specifically.

| EMDB id | CCC |  |
| --- | --- | --- |
|  | Initial | Final |
| emd_6640 | 0.74 | 0.88 |
| emd_10097 | 0.63 | 0.74 |
| emd_3488 | 0.73 | 0.86 |
| emd_10092 | 0.69 | 0.89 |
| emd_30127 | 0.77 | 0.83 |
| emd_9102 | 0.74 | 0.90 |
| emd_10800 | 0.75 | 0.84 |
| emd_8194 | 0.32 | 0.39 |
| emd_3947 | 0.66 | 0.70 |
| emd_20278 | 0.23 | 0.28 |
| emd_8743 | 0.15 | 0.23 |
| emd_20857 | 0.80 | 0.83 |
| emd_21430 | 0.75 | 0.82 |
| emd_21431 | 0.76 | 0.81 |
| emd_0707 | 0.80 | 0.87 |
| emd_7875 | 0.69 | 0.78 |
| emd_3145 | 0.17 | 0.23 |

**Extended Data Fig. S1.**
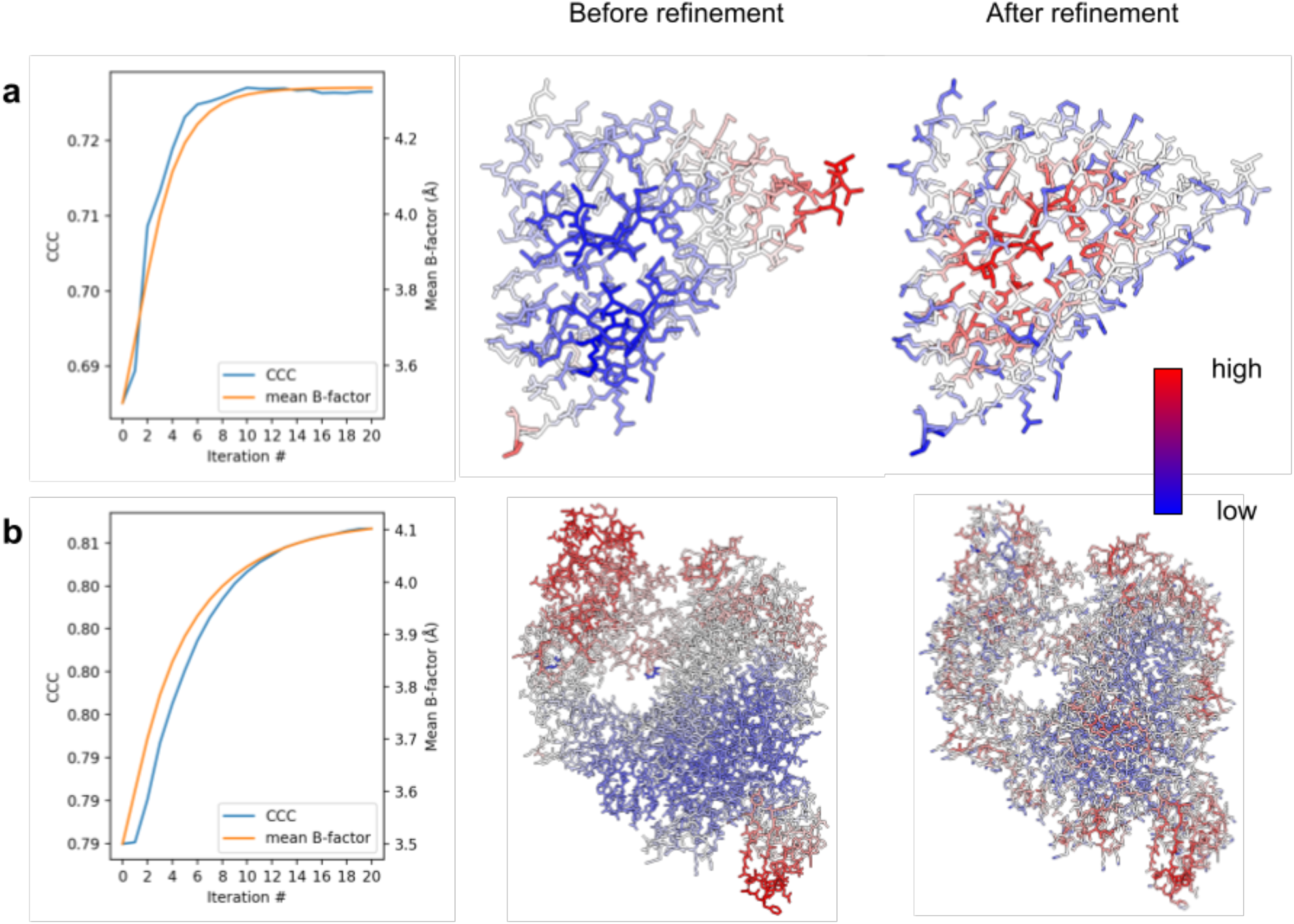
Change in CCC and B-factor during the TEMPy-REFF refinement procedure. **a,** CCC and B-factor changes illustrated on EMD-10097; **b,** and EMD-30127. The B-factor converges in around 10-12 iterations, thereby improving the CCC, i.e. the quality of fit to the data (left column). Significant changes in B-factor distribution can be seen before (middle column) and after refinement (right column).

**Extended Data Fig. S2.**
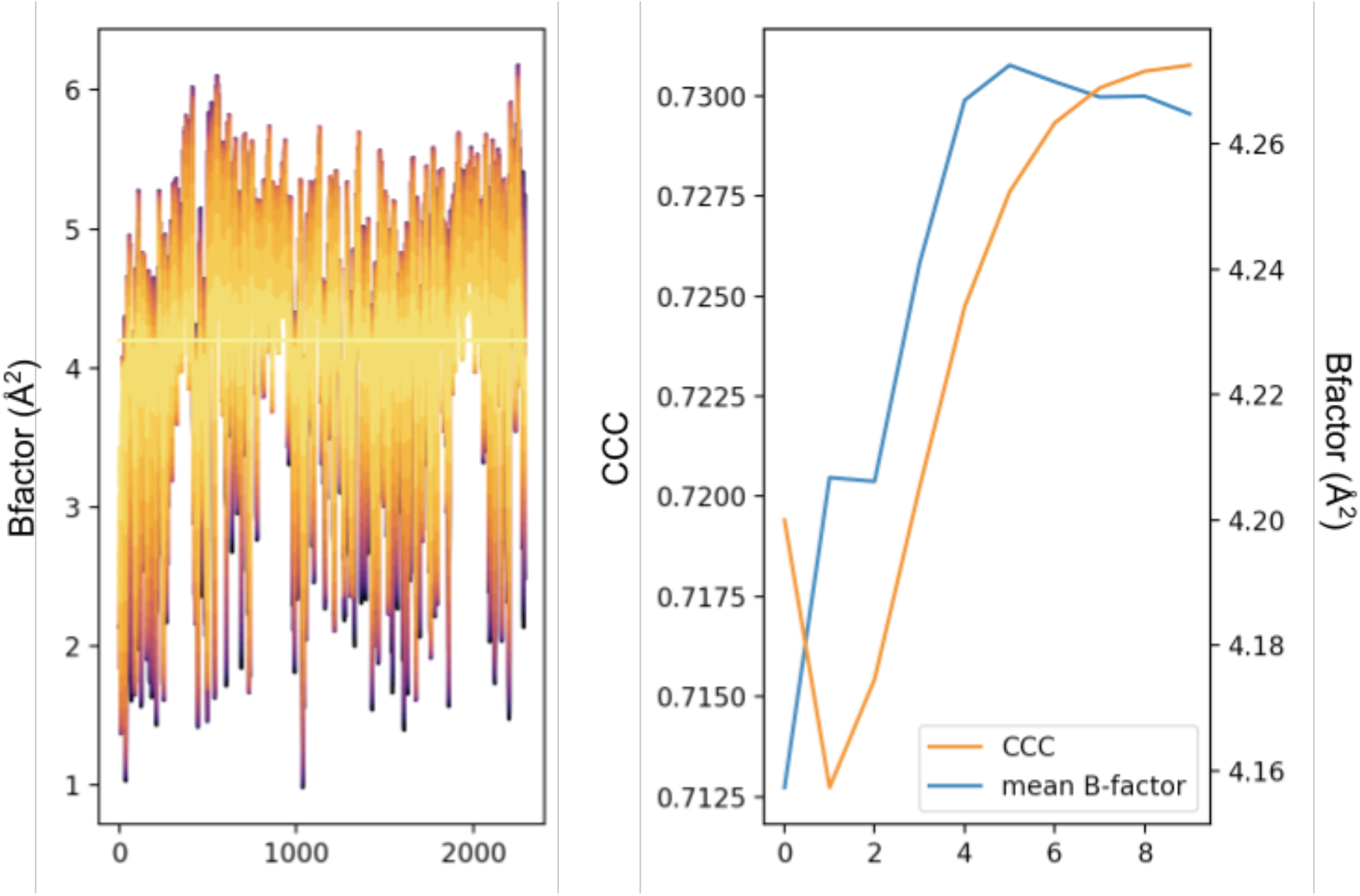
The change in B-factors during the TEMPy-REFF refinement process. Starting from an initial assignment, the B-factor converges to their final value, with an accompanying improvement in CCC.

**Extended Data Fig. S3.**
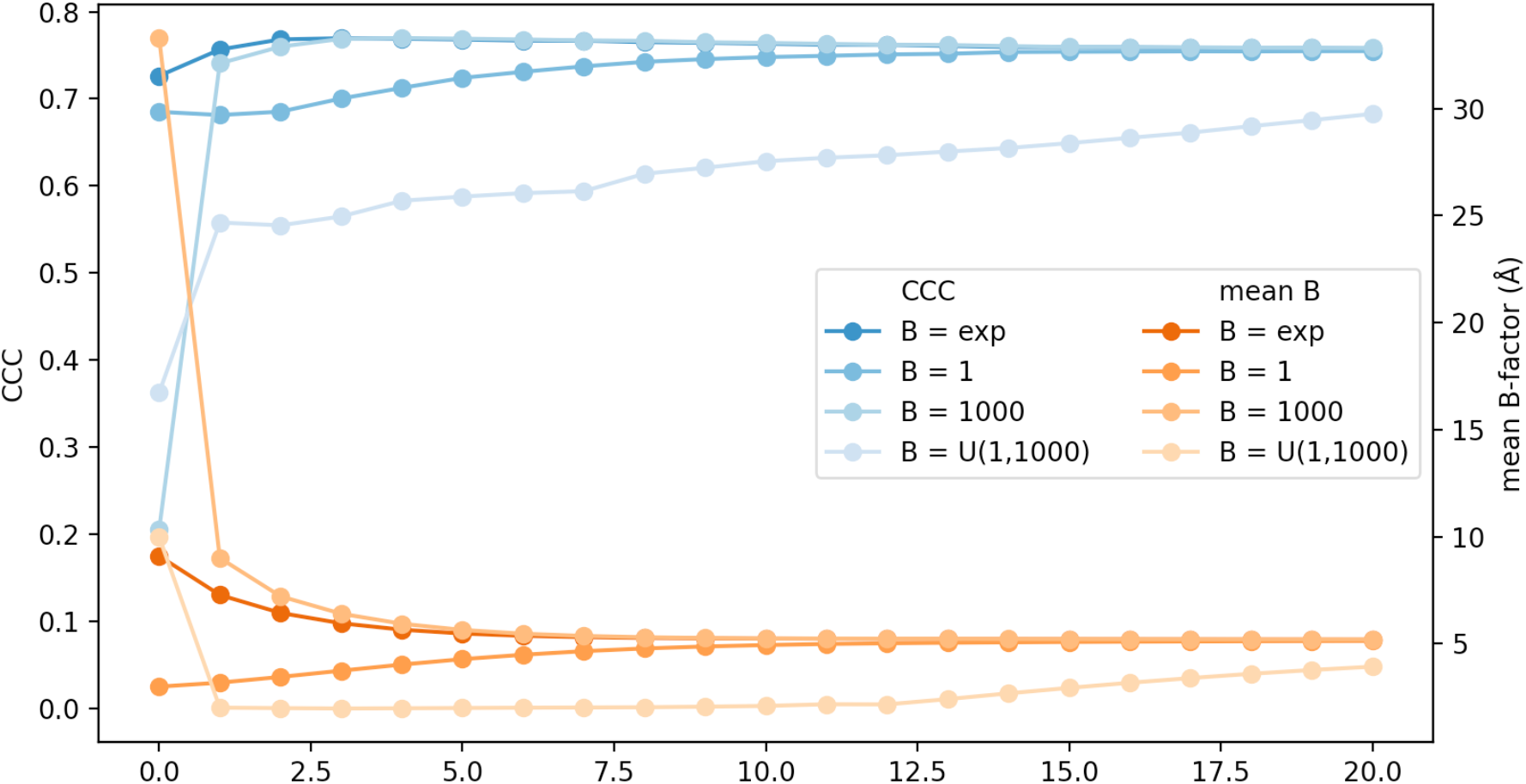
The convergence of B-factor values during the TEMPy-REFF refinement process, using multiple initial B-factor assignments: Unit B-factors to all atoms (all B = 1); All B = 1000; Random uniform B-factors between 1 and 1000 (B = U(1, 1000)); Random normal B-factors, with mean and variance according to the experimental distribution (if either of those value was unavailable, it was initialised to 1) (B = exp)

**Extended Data Fig. S4.**
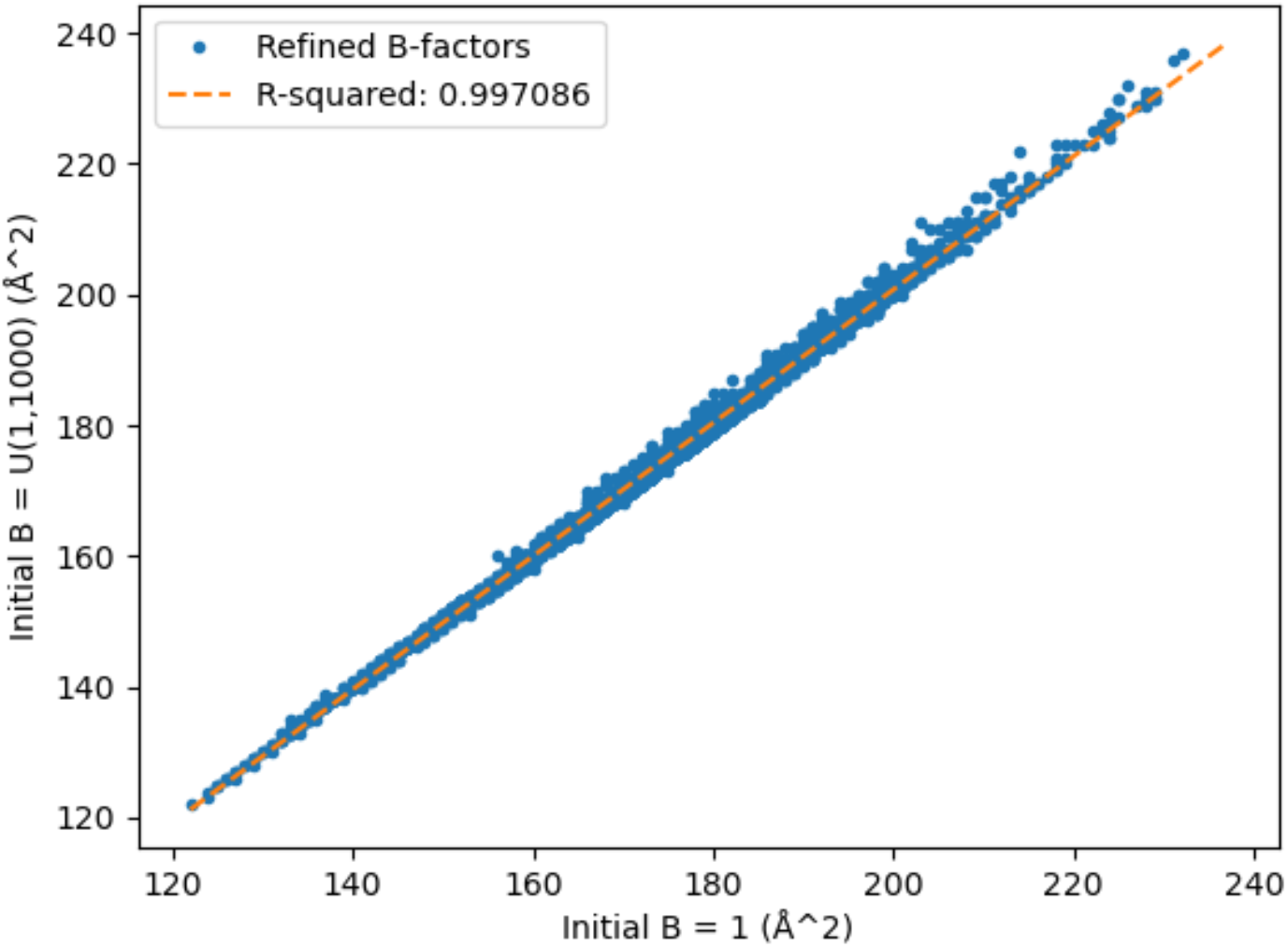
The values of the final B-factor after TEMPY-REFF refinement, for an initial uniform B-factor of 1 vs uniformly distributed between 1 and 1000. The B-factors are largely independent from the initial value.

**Extended Data Fig. S5.**
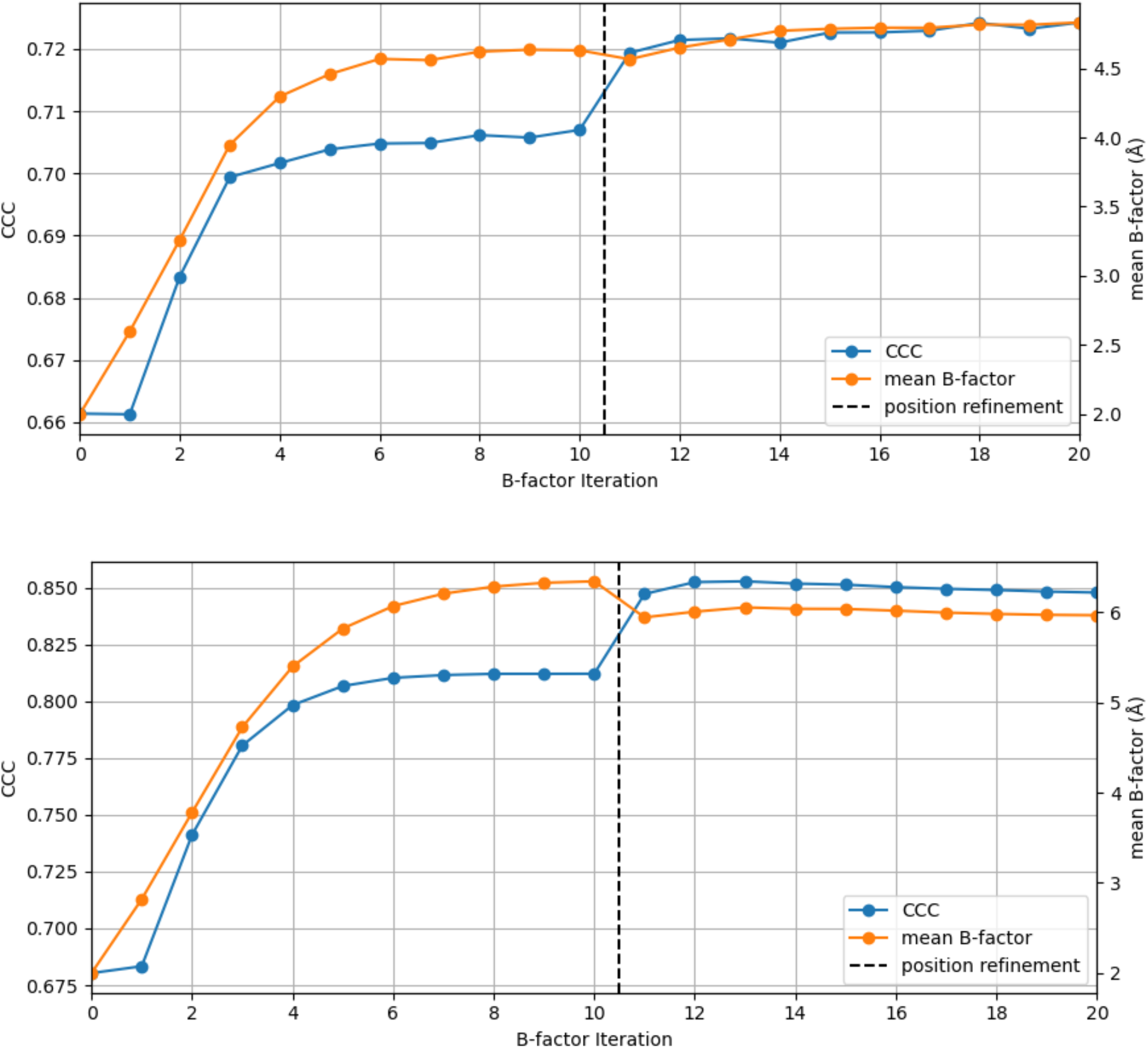
The change in Bfactor, before and after position refinement for two assemblies. **a,** EMD-0407; PDB ID 6NBC **b,** EMD-30127; PDB ID 6M71

**Extended Data Fig. S6.**
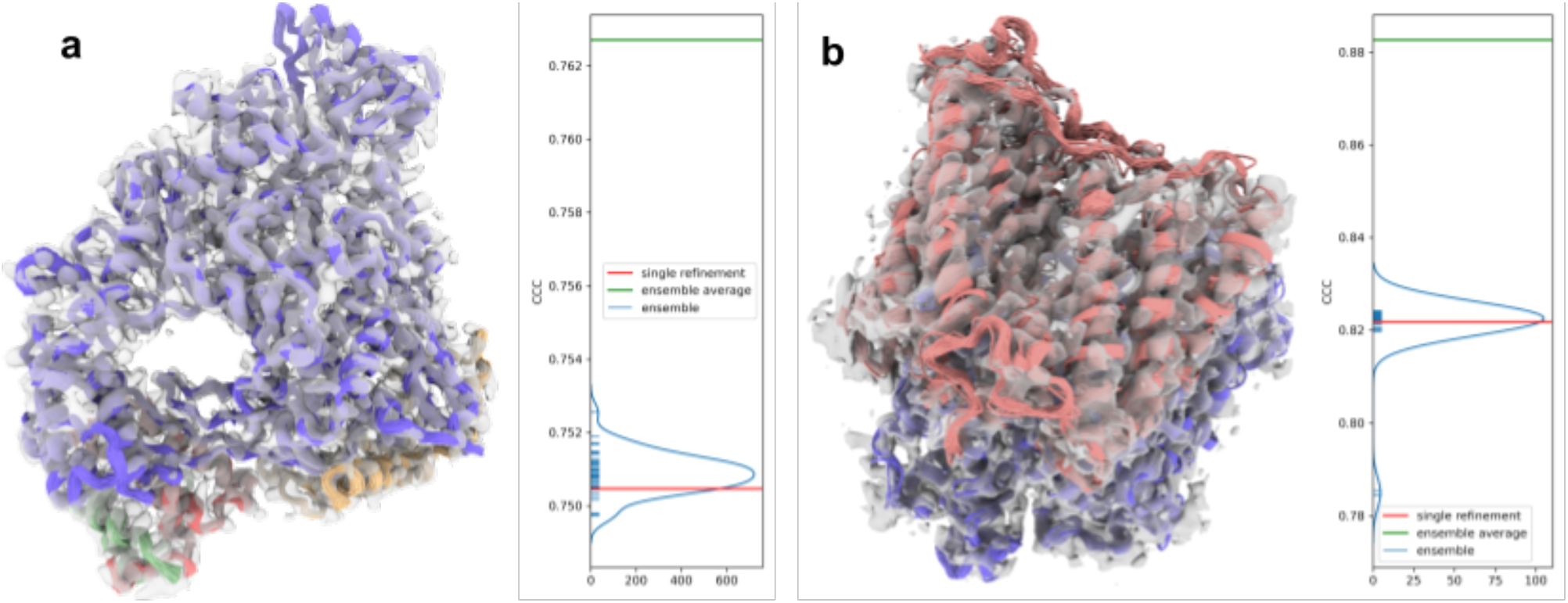
Ensemble generation. **a**, depiction of an ensemble of structures for SARS-Cov-2 RNA-dependent RNA polymerase at 2.9 Å resolution (EMD-30127, initial model PDB ID: 6M71). The CCC with respect to the map is shown on the right, with the single refinement shown in red, the ensemble in blue, and the ensemble average map in green. b) similar depiction for otopetrin proton channel Otop3 at 3.22 Å (EMD-9361, initial model PDB ID: 6NF6).

**Extended Data Fig. S7.**
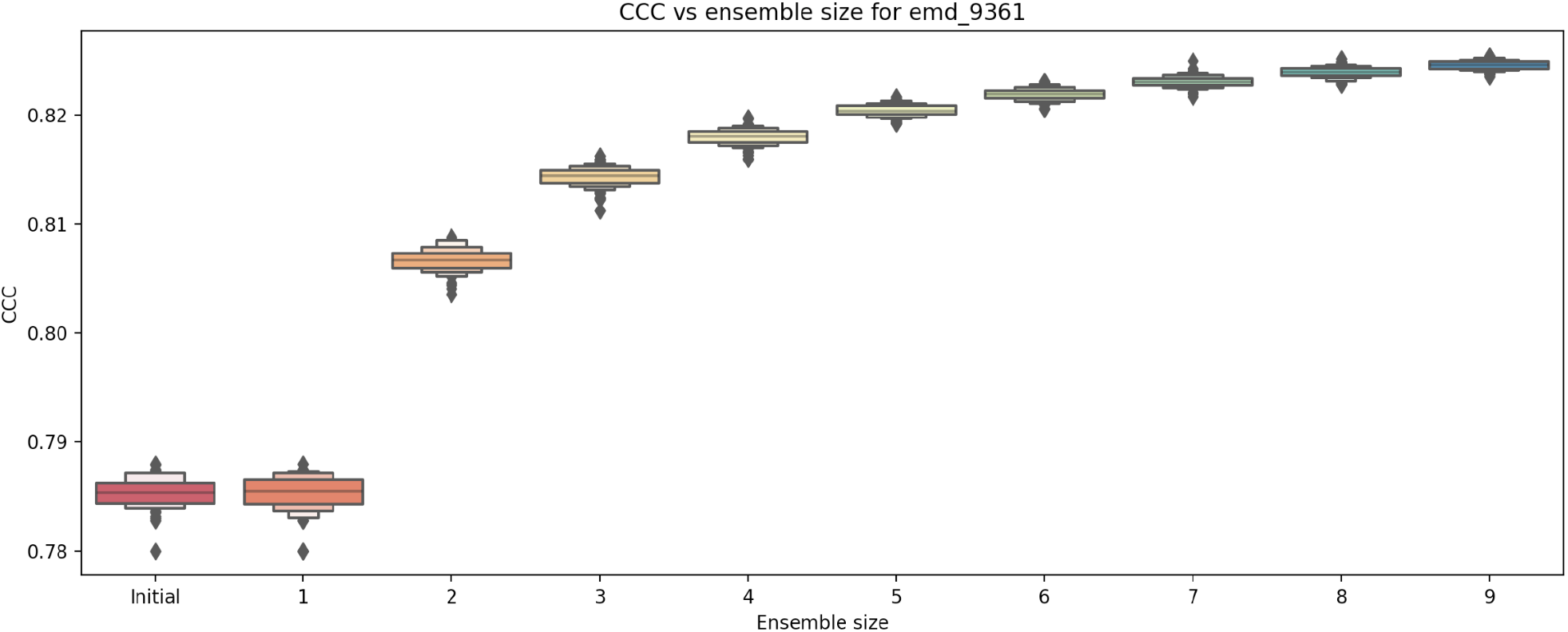
Change in CCC as a function of ensemble size, for the otopetrin proton channel Otop3 at 3.22 Å (EMD-9361, initial model PDB ID: 6NF6). Bootstrap estimates at each size are computed by resampling from the full ensemble.

**Extended Data Table S2.** Change in scores upon refinement, comparison against CERES.

| EMDB id | CCC |  |  | Clash score |  |  | Molprobity score |  |  | CaBLAM score |  |  |
| --- | --- | --- | --- | --- | --- | --- | --- | --- | --- | --- | --- | --- |
|  | Initial | Final | CERES | Initial | Final | CERES | Initial | Final | CERES | initial | Final | CERES |
| emd_0707 | 0.741 | 0.869 | 0.772 | 57.95 | 20.82 | 11.68 | 3.74 | 2.72 | 2.15 | 25.9 | 21.5 | 17.1 |
| emd_0709 | 0.628 | 0.913 | 0.870 | 69.25 | 21.73 | 14.30 | 3.79 | 3.11 | 2.20 | 24.6 | 23.4 | 15.9 |
| emd_10058 | 0.332 | 0, co | 0, co | 9.05 | 15.13 | 9.17 | 1.75 | 2.44 | 1.72 | 5.2 | 6.3 | 6.1 |
| emd_10092 | 0.748 | 0.925 | 0.733 | 15.84 | 25.56 | 11.08 | 2.45 | 2.72 | 2.04 | 20.9 | 14.8 | 17.0 |
| emd_10097 | 0.887 | 0.955 | -0.03 | 9.60 | 13.05 | 20.36 | 2.84 | 2.15 | 2.36 | 14.1 | 9.9 | 13.6 |
| emd_10658 | 0.610 | 0.884 | 0.754 | 100.71 | 24.27 | 5.58 | 3.81 | 2.84 | 1.30 | 14.8 | 16.8 | 3.5 |
| emd_20857 | 0.727 | 0.877 | 0.667 | 10.93 | 26.33 | 7.03 | 2.09 | 2.33 | 1.70 | 9.6 | 7.8 | 7.2 |
| emd_21247 | 0.710 | 0.799 | 0.657 | 23.91 | 23.72 | 7.62 | 2.74 | 2.41 | 2.15 | 16.2 | 13.9 | 13.2 |
| emd_21430 | 0.809 | 0.906 | 0.784 | 5.01 | 28.12 | 8.75 | 1.47 | 2.21 | 1.47 | 5.5 | 3.9 | 5.3 |
| emd_21431 | 0.786 | 0.917 | 0.757 | 4.81 | 29.41 | 8.01 | 1.25 | 2.28 | 1.59 | 6.9 | 4.4 | 5.1 |
| emd_30127 | 0.800 | 0.900 | 0.785 | 25.33 | 21.51 | 8.70 | 2.86 | 2.26 | 1.71 | 11.1 | 9.2 | 6.9 |
| emd_6272 | 0.903 | 0.982 | -0.3, | 22.33 | 26.88 | 8.93 | 2.48 | 2.41 | 1.62 | 4.3 | 7.1 | 5.6 |
| emd_6640 | 0.779 | 0.934 | 0.824 | 136.69 | 28.26 | 17.95 | 4.49 | 3.22 | 2.29 | 20.9 | 22.0 | 12.6 |
| emd_6904 | 0.533 | 0.865 | 0.855 | 83.30 | 19.51 | 28.37 | 4.22 | 3.23 | 2.60 | 35.8 | 30.9 | 17.3 |
| emd_9102 | 0, co | 0, co | 0, co | 83.18 | 19.47 | 19.43 | 3.71 | 3.24 | 2.27 | 9.7 | 19.1 | 9.8 |
| emd_9361 | 0.224 | 0.918 | 0.524 | 63.42 | 31.23 | 11.96 | 3.86 | 3.24 | 1.87 | 20.4 | 19.3 | 6.5 |

**Extended Data Table S3.** Speed benchmark. time for a run, for a given system

| EMDB id | Time |
| --- | --- |
| emd_0407 | 2m 20s 930ms |
| emd_0707 | 26m 57s 338ms |
| emd_0709 | 40m 24s 471ms |
| emd_10058 | 2s 579ms |
| emd_10092 | 11m 18s 315ms |
| emd_10097 | 55s 787ms |
| emd_10658 | 32m 20s 303ms |
| emd_10800 | 5m 6s 755ms |

**Extended Data Table S4.** Forcefield comparison: change in CCC after refinement using either AMBER orCHARMM. To confirm that no significant bias is introduced by the choice of forcefield, we tested both AMBER and CHARMM36 forcefields on our dataset. No significant differences are observed using either AMBER or CHARMM forcefields, although the CCC at the end of the refinement were slightly higher for the AMBER runs, on average (Runs with the highest CCC for a given system are highlighted in green).

| PDB | Initial CCC | Final CCC with Amber | Final CCC with CHARMM |
| --- | --- | --- | --- |
| 1a45 | 0.9306 | 0.962 | 0.9612 |
| 1ckm | 0.9019 | 0.9564 | 0.9553 |
| 1ffg | 0.9507 | 0.9607 | 0.9608 |
| 1hrd | 0.9218 | 0.964 | 0.9638 |
| 1ikn | 0.9121 | 0.9552 | 0.954 |

**Extended Data Fig. S8.**
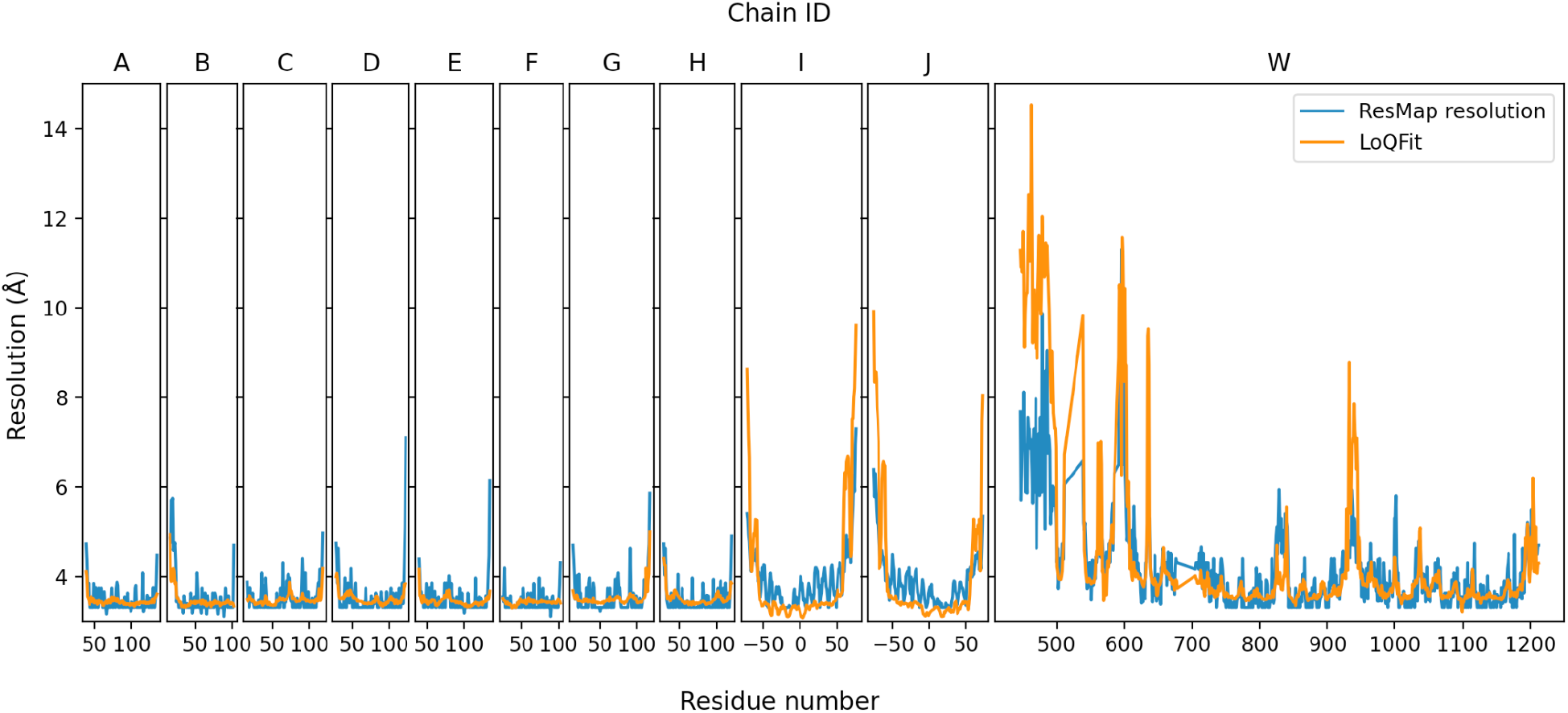
Comparison of LoQFit and ResMap local resolution for the nucleosome-CHD4 complex. Both follow very similar trends, with regions of locally higher and lower resolutions in agreement between the methods. (To obtain a local resolution along the chain for ResMap, we compute the average local resolution of the voxels in which the given residue is contained).

**Extended Data Fig. S9.**
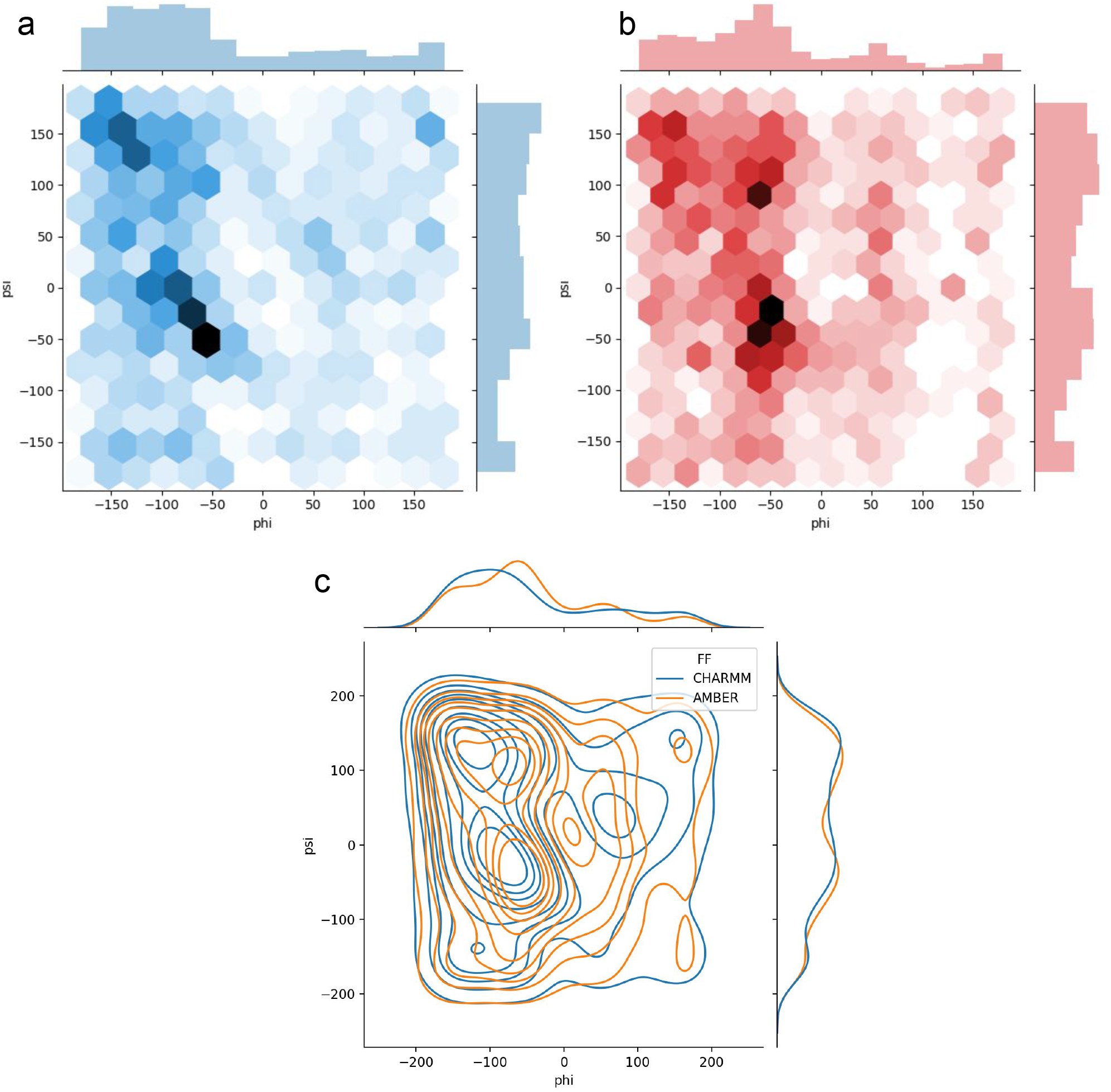
Ramachandran plot for **a,** CHARMM and **b,** AMBER: although similar, both forcefields exhibit slightly different preferences regarding dihedral combinations; the centers are slightly offset (most visible in **c,**), and differently populated.

**Extended Data Fig. S10.**
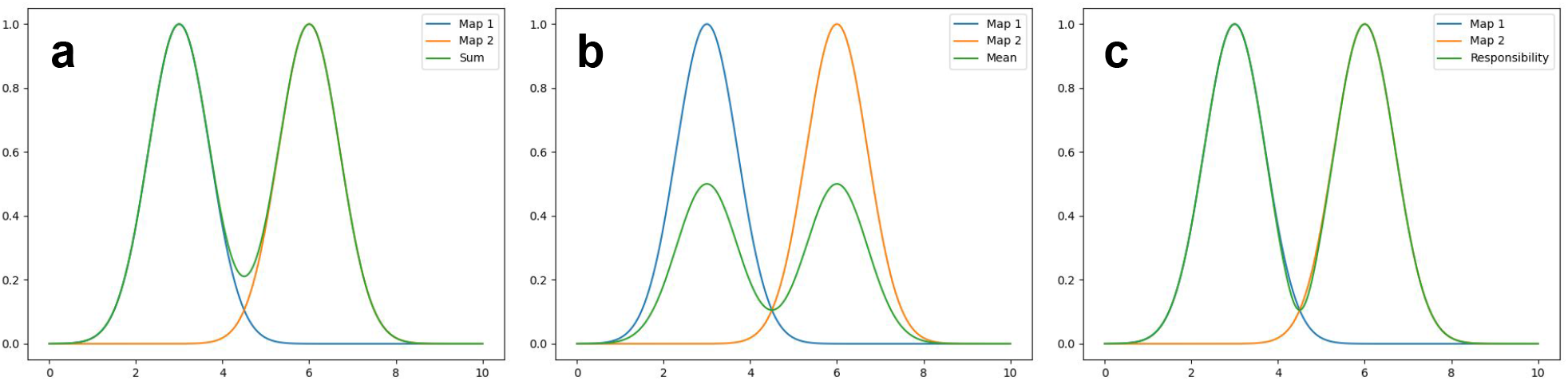
Schematics of map composition with responsibilities. We use two Gaussians to illustrate the map composition process (blue and yellow). Each Gaussian is an idealised representation of a focused map. Three protocols for combining those maps into a single composed map are shown, with the resulting map in green. **a,** Summing the intensities from multiple maps will overcount the transition regions (creating a ‘seam’). **b,** Averaging the maps will undercount the peak intensities **c,** Using responsibilities, the weighting of each map switches smoothly from map1, to equal weighting at the seam, to map2. This results in properly accounting for the intensities across the whole domain.

